# Shifts in the thermal niche of fruit trees under climate change: the case of peach cultivation in France

**DOI:** 10.1101/2020.09.28.315960

**Authors:** C. Vanalli, R. Casagrandi, M. Gatto, D. Bevacqua

**Affiliations:** Center for Infectious Disease Dynamics and Department of Biology, The Pennsylvania State University, University Park, 16082 PA, USA; Dipartimento di Elettronica, Informazione e Bioingegneria, Politecnico di Milano, 20133 Milano, Italy; Plantes et Système de cultures Horticoles (PSH), UR 1115, INRAE, 84000, Avignon, France

**Keywords:** Process-based suitability model, global warming, blooming time, plant dormancy, peach (*Prunus persica*), plant phenology

## Abstract

Climate influences plant phenological traits, thus playing a key role in defining the geographical range of crops. Foreseeing the impact of climate change on fruit trees is essential to inform policy decisions to guide the adaptation to new climatic conditions. To this end, we propose and use a phenological process-based model to assess the impacts of climate change upon the phenology, the suitability and the distribution of economically important cultivars of peach (*Prunus persica*), across the entire continental France. The model combines temperature dependent sub-models of dormancy, blooming, fruit survival and ripening, using chilling units, forcing units, frost occurrence and growing degree days, respectively. We find that climate change will have divergent impacts upon peach production. On the one hand, blooming will occur earlier, warmer temperatures will decrease spring frost occurrence and fruit ripening will be easily achieved before the start of fall. On the other hand, milder winters will impede the plant buds from breaking endodormancy, with consequent abnormal patterns of fruit development or even blooming failure. This latter impact will dramatically shift the geographic range of sites where peach production will be profitable. This shift will mainly be from the south of France (Languedoc-Roussillon, Rhône-Alpes and Provence-Alpes-Côte d’Azur), to northwestern areas where the winter chilling requirement will still be fulfilled. Our study provides novel insights for understanding and forecasting climate change impacts on peach phenology and it is the first framework that maps the ecological thermal niche of peach at national level.

## 1 Introduction

Climate plays a key role in defining the geographic range of plants (Whittaker, 1975) and climate change is expected to severely influence plant distributions in the forthcoming decades (Lenoir et al., 2008; Morin et al., 2008; Chuine, 2010; Gritti et al., 2013; Zhao et al., 2018). Plant phenology is strongly responsive to temperature and, indeed, phenological changes have been among the first documented fingerprints of climate change (Menzel & Fabian, 1999; Körner & Basler, 2010; Lee et al., 2013; Wolkovich et al., 2017). Climate change will therefore have an impact on agricultural production by altering the geographical distribution of economically important crops (Tao et al., 2006; Duchêne et al., 2010; Teixeira et al., 2011; Ghrab et al., 2014). Foreseeing this impact is essential to alert stake holders, inform decisions to implement adaptation strategies and to alleviate damaging consequences. Although some of the impacts can be mitigated in agricultural settings (e.g. irrigation, frost protection), the challenge will be greater for perennial crops. These are subject to climate impacts throughout the year and their decades-long lifespan makes the choice of where to plant an orchard critical (Lobell & Field, 2012).

To date, spatial shifts in the distribution of crops have been assessed using empirical Species Distribution Models (SDMs) (Machovina & Feeley, 2013), as well as process-based Suitability Models (SMs) (e.g. White et al., 2006; Keenan et al., 2011). Both modeling frameworks have merits and shortcomings. However, as noticed by Parker and Abatzoglou (2017), “unlike SDMs, SMs can provide information on specific climatic limitations and crop phenology, and are not limited by the correlative approach”. When dealing with perennial plant phenology and climate, the blooming process is probably the most studied. Until one decade ago, most studies provided consensus on the earlier occurrence of plant blooming. For example, Estrella et al. (2007) reported that phenological events such as emergence and blooming “are significantly earlier now than 53 years ago, with a mean advance of 1.1-1.3 days per decade”. According to Chmielewski et al. (2004) “phenological phases of the natural vegetation as well as of fruit trees and field crops have advanced clearly in the last decade of the 20^th^ century”. Nevertheless, in the last years some authors theorized a possible trend reversal due to a subtle, process-dependent cause: strong warming in winter could slow the fulfillment of chilling requirements, which may delay spring phenology (Hänninen & Tanino, 2011). These scenarios were also confirmed in recent field studies (Yu et al., 2010; Laube et al., 2014). Accordingly, models that explicitly considered the fulfillment of both chilling and forcing requirements, also called two-phase or sequential Chilling/Forcing (CF) models, were proposed to better assess the date of blooming. For the genus *Prunus*, this has been accomplished for apricot and peach trees (Chuine et al., 2016) as well for cherry trees (Chmielewski & Götz, 2016).

To design adaptation strategies, the occurrence of blooming is a necessary yet not sufficient condition to make an area suitable for fruit production. An area is considered suitable if the environmental conditions allow all the processes leading to full fruit ripeness to occur. Following this rationale, Parker and Abatzoglou (2017) assessed possible shifts in the thermal niche of almond trees under climate change in the Western United States. Santos et al. (2016) assessed values of chilling and heat accumulation over Portugal and discussed possible related shifts of the thermal niche of several fruit classes (from carob to lemons and from olives to vines). Similarly, Ahmadi and Baaghideh (2018) assessed the impact of climate change on apple tree cultivation in Iran. They assumed that a given area would be suitable if temperatures remained within certain boundaries in any plant development stage.

Here, we propose a novel modeling approach that combines temperature-dependent phenological models of blooming, fruit survival and ripening to assess suitability for peach *Prunus persica* tree cultivation. We optimize the models for nine different peach cultivars and demonstrate our approach for a reference past period (1996-2015). Then, we use it to project shifts in the peach cultivation range in continental France under different scenarios, that are Representative Concentration Pathways, RCPs 4.5 and 8.5 (IPCC, 2013) of average and minimum daily temperature change in the near (2021-2040) and far (2081-2100) future. Globally, the peach is the third most cultivated plant of the *Rosaceae* family (Obi et al., 2018). This fruit has been extensively studied in both field and modeling works (Génard & Huguet, 1996; Ziosi et al., 2003; Allen et al., 2006; Miras-Avalos et al., 2011) and its sensitivity to climate change has already been established (Litschmann et al., 2008; Ghrab et al., 2014). The French territory is a paradigmatic case for studying peach cultivation, because it covers four climatic zones (Mediterranean, continental, oceanic and mountain) in less than 600,000 km^2^.

## 2 Materials and Methods

### 2.1 Phenological dates

From the database TEMPO (National Network of Phenology Observatories) (INRA, Seguin, 2004), we obtained 159 pairs of data that report blooming and harvest dates for nine peach cultivars (Snowqueen, M. Sundance, Springlady, OHenry, Alexandra, Flavorglod, Benedicte, Flavorcrest and Emeraude) in the period 1987-2008. These concern three experimental sites in southern France: Bordeaux (long. 0° 34′ W, lat. 44° 46′ N), Balandran (long. 4° 28′ E, lat. 43° 45′ N) and Étoile-sur-Rhône (long 4° 53′ E, lat. 44° 49′ N) (see Supplementary Information Figure S1).

### 2.2 Temperature: records, assessments and projections

Daily average temperature data (1987-2008) were obtained from meteorological stations in the proximity of the experimental sites (see Supplementary Information Table S1) and provided by the INRA CLIMATIK platform (https://intranet.inra.fr/climatik_v2). We obtained assess-ments of daily minimum and average temperatures for the entire French continental area for the period 1996-2015. Assessments were generated by the EUROCORDEX CNRM-CERFACS CM5 model and downscaled with a spatial resolution of around 8 × 8 km^2^ (~ 0.11° × 0.11°). They have been made available by the project Drias developed by Météo-France (Drias, 2013). The same project provided projections of the daily minimum and average temperature for two Representative Concentration Pathways scenarios, RCP 4.5 and RCP 8.5 (IPCC, 2013), in the period 2020-2100, downscaled at the same spatial resolution, for the entire French territory. The RCPs refer to different emission scenarios providing an estimated increase of global mean surface temperatures at the end of the 21^st^ century, is likely to be in the ranges of 1.1-2.6°C for RCP 4.5 and 2.6-4.8°C for RCP 8.5. To analyze the extent of climate warming in France, we selected four cells of 24 × 24 km^2^, one for each climatic zone (Mediterranean, continental, oceanic and mountain), and we evaluated the daily temperature anomalies in the the near (2021-2040) and far (2081-2100) future with respect to a reference time period (1996-2015).

### 2.3 A model to predict blooming date

First, we performed a 1-way ANalysis Of VAriance (ANOVA, with a significance level of *α* = 0.05) to test if observed blooming dates significantly differed between the nine considered cultivars. Second, we used the two-phases model proposed by Chuine et al. (2016) to predict blooming dates. That model assumes that endodormancy break occurs at the time *t_C_*, when the state of chilling *S_c_*(*t*), resulting from the sum of the daily rates of chilling *R_c_* (as detailed below, see eq. 5), reaches the critical value *C*\*. According to Chuine et al. (2016), thus, we compute

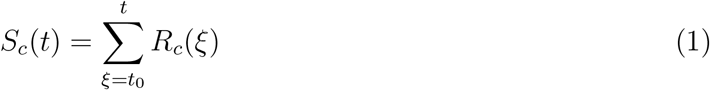

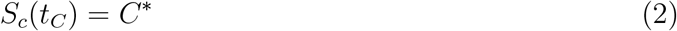

with *t* measured in days and *t*_0_ fixed to September 1^st^. Blooming is assumed to occur at time *t_B_* when the state of forcing *S_f_*, i. e. the sum of the daily rates of forcing *R_f_* (also detailed below, see eq. 6), reaches the critical value *F*\*. According to Chuine et al. (2016), we have

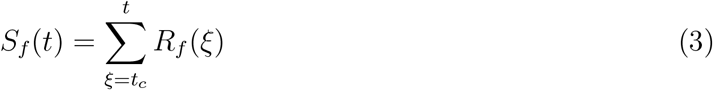

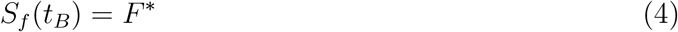

with *t_C_* determined by equation 2. Note that the endodormancy break is a necessary condition to enter the ecodormancy phase that precedes blooming. Both daily rates of chilling *R_c_* and forcing *R_f_* are functions of the average daily temperature *T*(*t*), itself varying with time t. We computed *R_c_* using a symmetrical and unimodal function proposed by Chuine (2000):

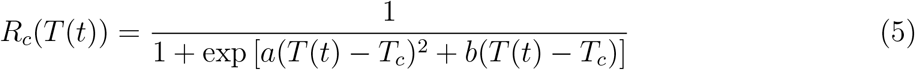

where *a* (in units of °C^−2^), *b* (°C^−1^) and *T_c_* (°C) are species-specific parameters that describe the accumulation of chilling units. According to Chuine et al. (2016), we computed *R_f_* as a logistic function of temperature:

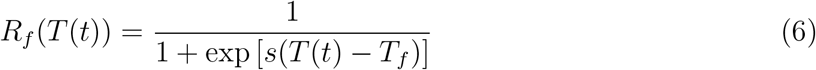

where the parameter *s* (°C^−1^) shapes the steepness of the curve and *T_f_* (°C) the logistic midpoint. As for the chilling function, we used the parameters optimized by Chuine et al. (2016) for peach (*a* = 3.53, *b* = −25.85, *T_c_* = 1.52, *C*\* = 49.6), while we optimized the parameters of the forcing function, *i. e. s, T_f_* and the required *F*\*, using our data set by minimizing the Root Mean Square Error (*RMSE*, see Burnham and Anderson, 2002):

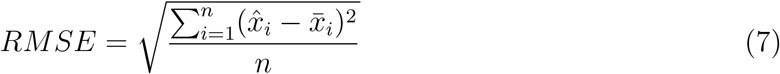

where 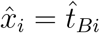 is the estimated blooming date, 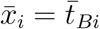 is the observed one, and *n* the number of observations over three different sites. We performed the *RMSE* minimization via the Nelder-Mead simplex algorithm (Lagarias et al., 1998), using the built-in MATLAB optimizer “fminsearch”.

### 2.4 A model to predict ripening duration

We performed a 1-way ANalysis Of VAriance (ANOVA, with a significance level of *α* = 0.05) to test if the observed ripening duration *d_R_* (i. e. the time from blooming = *t_B_* to ripening = *t_R_*) was significantly affected by the factor “cultivar”. In case of significance, we performed a multiple t-test using Scheffé’s procedure (Savin, 1980), with a significance level *α* = 0.05, to evaluate if cultivars could be classified in groups. If this was the case, we calibrated a different parameter set of the ripening model for each group (see below). We assume that a fruit is ripe when the state of ripening *S_R_, i.e*. the sum of the daily rates of heating *R_R_* (see eq. 10 below), reaches the critical value *R*\* (Miller et al., 2001; Kenealy et al., 2015):

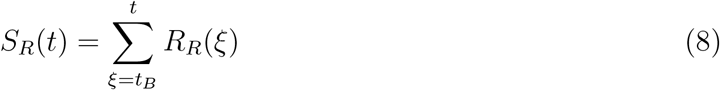

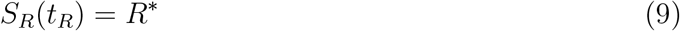

where *t_B_* is the observed blooming time for each cultivar and year. *R_R_*(*t*) is calculated as

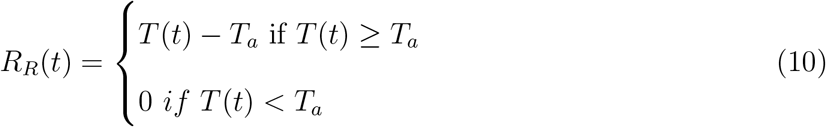

where *T_a_* is the base activation temperature, which we set equal to 7°C (Miller et al., 2001; Kenealy et al., 2015). Thus, R* is the sum of Growing Degree Days (GDD). Then, for each cultivar group we estimated the value of R* minimizing the *RMSE* (see eq. 7), where 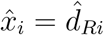 is the estimated ripening duration, 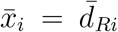 is the observed one, and *n* is the number of observations. If ripening is not achieved by the 21^st^ of September, when the plant is assumed to enter dormancy (Battey, 2000; Gauzere et al., 2017), the yield is considered to be entirely lost. In other words, the environmental conditions are not suitable for cultivation.

### 2.5 Thermal niche for fruit tree cultivation

We assume that an area is suitable for peach cultivar if all three necessary conditions are sequentially met: i) blooming can occur, i. e. both chilling and forcing requirements are satisfied; ii) no frost events occur at the turn of blooming; and iii) ripening can occur. To model the fact that flowers (considered here as newborn fruits) are sensitive to frost in their first week after blossom (Rodrigo, 2000), we assume that fruits do not survive if there is at least one day, in the period from *t_B_* − 3 to *t_B_* + 3, with minimum daily temperature below a critical temperature (*T_x_*). For peach, according to Snyder and Melo-Abreu (2005), this critical temperature corresponds to −4.9 °C. We assumed that orchards can be irrigated so that water requirements are met independently from climatic conditions. We assessed the spatial suitability for different varieties of peach cultivation across France (8412 map cells of 8×8 km^2^) over three different periods, i.e. a reference (1996-2015), a near future (2031-2050) and a far future (2081-2100) period.

We considered a map cell as suited for cultivation if all three conditions described above are accomplished for at least 18 years out of 20.

## 3 Results

### 3.1 Models for blooming and ripening date

We found that the factor “cultivar” has no significant effect on the observed blooming dates (*p* = 0.61). The values of parameters of the blooming model optimized against our data set, namely *s, T_f_* and *F*\* (see eq. 5), are −0.5 °C^−1^, 14.4 °C and 4.19, respectively. Although the model is quite hyperactive, *i.e*. it tends to underestimate earliest blooming dates and to overestimate the latest ones, it reproduces the observed data well (Figure 1a). We evaluated if the model residuals are cultivar-dependent via an ANOVA and the resulting p-value of 0.65 suggests that the model simulates the peach blooming time in an equivalent way for all the different cultivars.

**Figure 1:**
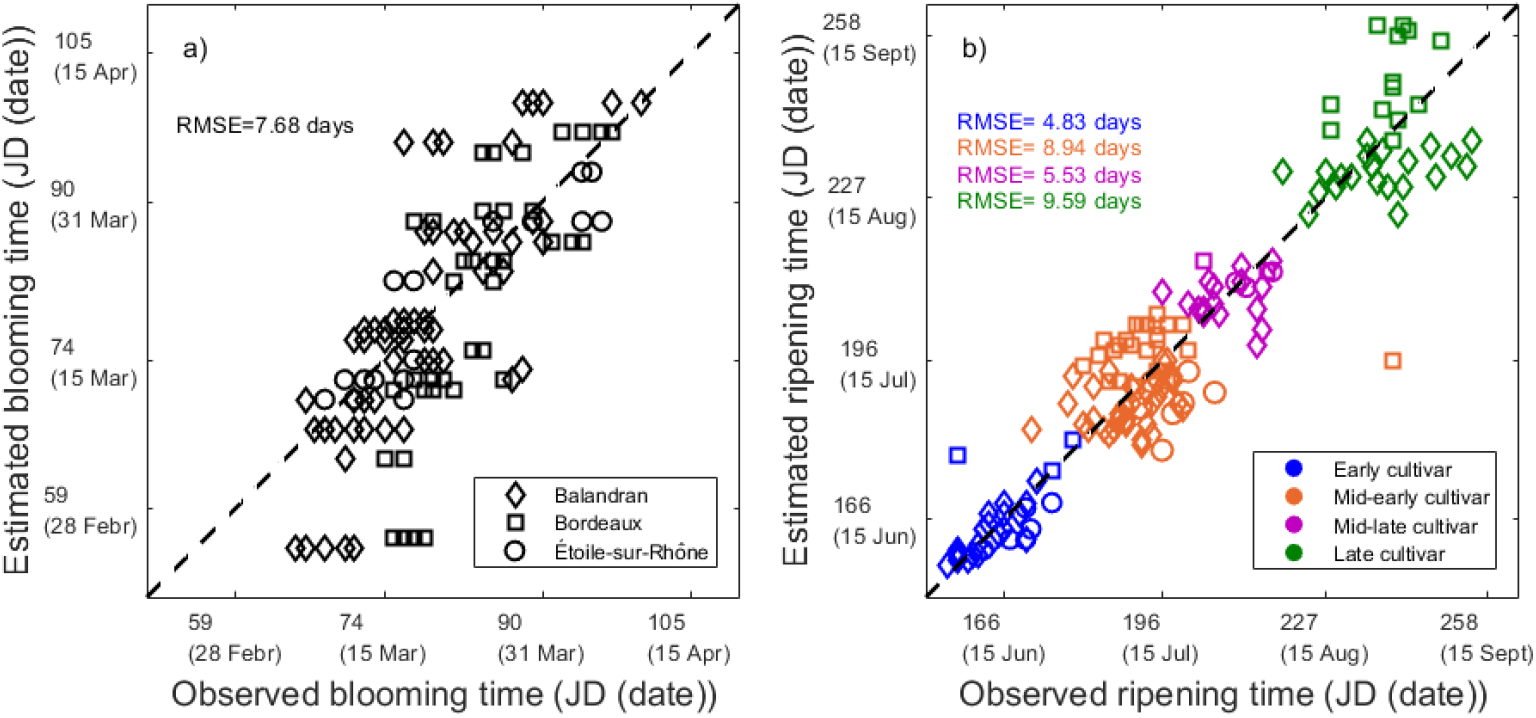
Blooming and ripening dates. Observed (x-axis) vs predicted (y-axis) dates in Julian Day (JD) and in calendar dates (within brackets) of (caption a) blooming and (caption b) ripening. Diamonds, squares and circles refer to the sites of Balandran, Bordeaux and Étoile-sur-Rhône, respectively. Ripening date is represented in blue for early, in orange for mid-early, in purple for mid-late and in green for late cultivars.

On the contrary, we interestingly found that the factor cultivar has a significant effect on the observed ripening duration (*p* < 0.01). We therefore refined the analysis for the ripening duration applying Scheffé’s procedure and performing a multiple t-test (Figure S2). The procedure identified four cultivar clusters that we can name: “early cultivar” (Alexandra and Springlady), “mid-early cultivar” (Snowqueen, Flavorgold, Flavorcrest and Emeraude), “mid-late cultivar” (Benedicte) and “late cultivar” (M.Sundance and OHenry). We then optimized the parameter of forcing requirement Growing Degree Days for each cultivar clustering, obtaining a GDD value equal to 678, 1026, 1371 and 1772, respectively, for the “early”, “mid-early”, “mid-late” and “late” cultivars. Adherence of model predictions (y-axis) to observations (x-axis), using our model with calibrated parameters for each cluster, is quite satisfactory, as shown in Figure 1b. We also evaluated how the error in the estimate of blooming time propagates to the ripening period duration. We used the estimated blooming time as initial time for the ripening time model and we compared the estimated ripening period duration with the observed data (Figure S3). Our analysis reveals that the predicted date of ripening does not vary significantly whether we used the described model (estimated) or the observed blooming date (early cultivar *p*= 0.95, mid-early cultivar *p*= 0.44, mid-late cultivar *p*= 0.24 and late cultivar *p*= 0.87).

### 3.2 Peach thermal niche in the reference period 1996-2015

Average temperature conditions over the reference period for the four considered cells representative of climatic zones are reported in the Supporting Information (Figure S4a-d and Table S2). Spring temperatures are warmer in the Mediterranean region [12-15 °C], with respect to those in the continental and oceanic regions [10-12 °C], which are comparable between them, while they are lower in the mountain areas [4-5 ^°^C]. Similarly, the warmest summers were registered in the Mediterranean region [20-22 °C], the coldest ones in the mountain areas [12-16 °C] and intermediate summers were found in oceanic and continental regions, where temperatures are more variable. Autumns and winters were characterized by milder temperatures in the Mediterranean [9-11 °C and 3-7 °C] and oceanic regions [8-12 °C and 4-7 °C].

As mapped in Figure 2a, our process-based suitability model estimates that for the reference period 1996-2015 the chilling requirement was achieved first (by the end of December) in the mountain regions (Alps, Pyrenees and in the Massif Central) and last (by mid February) in the Mediterranean and the Southern Atlantic (MSA) regions. Also, in a few map cells of these regions (notably, nearby the cities of Montpellier, Touloun, Sain Tropez, Perpignan et Bayonne) some winters were so mild that the chilling requirement was not satisfied and the endodormancy break was compromised. In contrast, blooming time (Figure 2b) occurred late (by end of April) in the mountain regions and first (by mid March) in the MSA regions. This occurs because in mountain regions, although the plants achieve chilling requirement first, they experience high temperatures triggering forcing rates much later in the season. Likewise, in some cells of mountain regions (Prealps and Massif Central), blooming was not effective because newborn fruits were injured by frost. Note that this was not the case for map cells at the highest altitudes were blooming occurred late enough to avoid frosting days. When considering the date of ripening (Figure 2c), a mid-early cultivar, which is expected to ripe around mid July in the MSA regions, has no time to ripe in the mountain regions. In the northern part of the country, instead, ripeness is achieved only at the end of August. Such a difference in the date of ripening is mostly due to blooming occurring later, rather than to lower summer temperatures. Mid-late and late cultivars need significantly more days to attain ripeness, so they can only be cultivated in those regions where blooming occurs first (Figure S5). For this reason, in terms of peach suitability (Figure 2d), the MSA regions were the most suited for peach cultivation, as both early to late cultivars could be grown and produce peach from mid June to the end of August. On the other hand, the mountain regions (Pyrenees, Massif Central and Alps, black in Figure 2d) were the only regions where no peach variety could be cultivated due to spring frost damages, late blooming and the consequent lack of time to attain ripeness.

**Figure 2:**
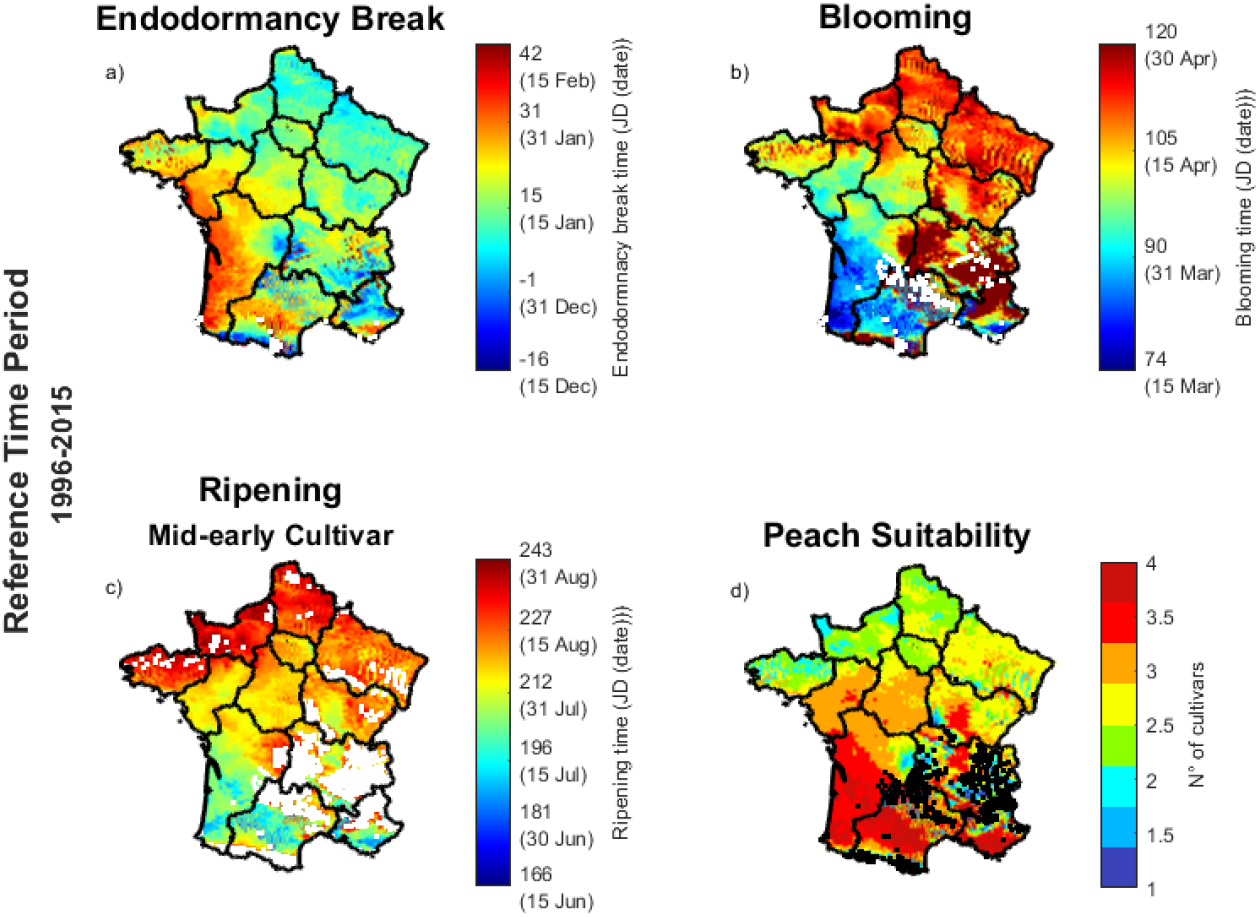
Peach phenological times and thermal niche for cultivation: hindcast in the French continental regions for the reference period (1996-2015). (a) Endodormancy break, (b) blooming, (c) ripening of mid early cultivars dates, in Julian Days and calendar dates (within brackets), and (d) peach suitability. For each map cell (8×8 km^2^), the average value over the considered 20 years is reported. White map cells (in captions a, b and c) represent those areas where the phenological event did not occur for at least two years. Likewise, black cells (in caption d) represent those areas where no peach cultivar could be cultivated for at least two years (out of 20).

### 3.3 Peach thermal niche in the future

According to the used scenarios (Figure S4 and Table S2), climate warming is expected to be more severe in the Mediterranean and mountain regions. Particularly, with the highest anomalies expected in winters for the Mediterranean regions, and in both summer and winters for the mountain areas. In the near future (2021-2040), the RCP 4.5 and RCP 8.5 scenarios are characterized by similar temperature anomalies. It is worthy to note that, for that period higher winter anomalies are generally expected for RCP 4.5 rather than for RCP 8.5. Conversely, in the far future (2081-2100), RCP 8.5 anomalies are expected to be consistently higher throughout the year, with the highest predicted warming in the mountain areas in summer (up to +5.1 °C).

Our mapping in Figure 3 reveals that the most suitable areas for peach cultivation in France are expected to significantly change in the near future (Figure 3a-d) and even more at the end of the century (Figure 3e-h). Some of the historically suitable zones in the MSA regions are predicted to become unsuitable because of blooming failure, being the chilling requirement impossible to achieve (Figure 3a,e). In those areas where the chilling requirement will still be satisfied, endodormancy break will be delayed. However, this will not directly reflect into a delayed blooming time, as the forcing requirement will be achieved more quickly. This earlier achievement of the forcing requirement will cause blooming time to occur 7-17 days in advance (Figure 3b,f). In other words, blooming is expected either to be impaired or to occur earlier. As a consequence of early blooming and warmer springs and summers, there will be an earlier occurrence of the ripening date (Figure 3d,g). In comparison with the reference period, the failures caused by not meeting the chilling requirement will cause numerous areas that are currently productive to become unsuitable. This is expected mainly in the Mediterranean regions and will be an important issue in the far future (see Figure 3e), especially for the RCP 8.5 scenario (see Figure S6e). In contrast, spring frost events and ripening failures will decrease, making continental areas in the northern part of the country more suitable for peach cultivation. Changes will be exacerbated in the far future of the RCP 8.5 scenario (Figure S6). In this far future scenario, there will be a paradoxical geographical divide, whereby the most suitable areas will be close to those that will not be able to host any cultivar because of the inability to bloom, while conditions would be optimal for fruit survival (i. e. no frost) and ripening.

**Figure 3:**
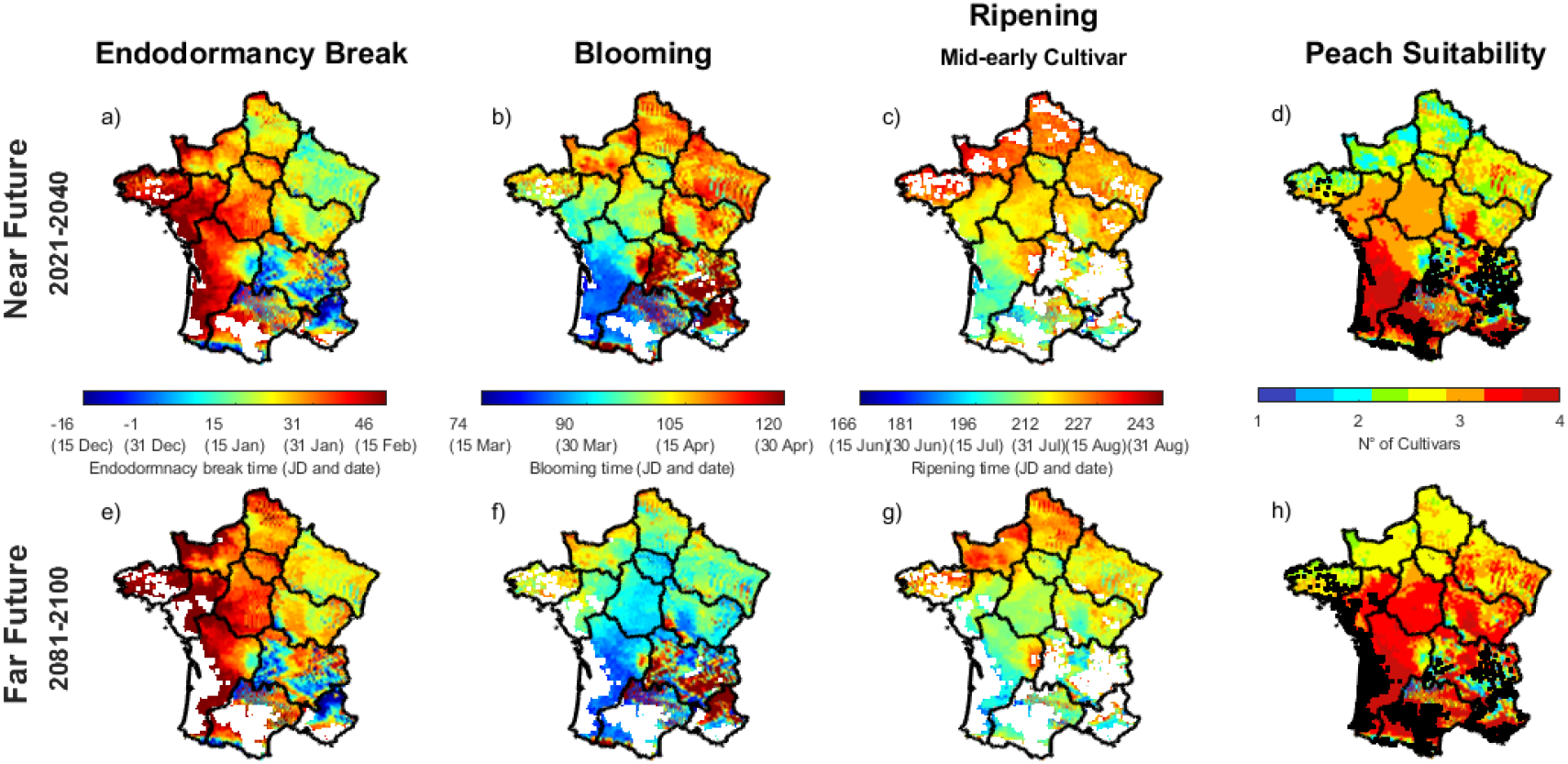
Peach phenological times and thermal niche for cultivation: scenarios forcing our model with RCP 4.5 in the French continental regions for the near (2021-2040, upper panels) and far (2081-2100, lower panels) future. (a,e) Endodormancy break, (b,f) blooming, (c,g) ripening of the mid-early cultivars dates, in Julian Days (JD) and calendar dates (within brackets), and (d,h) peach suitability. For each map cell (8×8 km^2^), the average value over the considered 20 years is reported. White map cells (a-c and e-g) represent those areas where the phenological event did not occur for at least two years. Likewise, black cells (d and h) represent those areas where no peach cultivar could be cultivated for at least two years (out of 20).

## 4 Discussion

### 4.1 Peach thermal niche in the reference period 1996-2015

Chill accumulation has historically not been considered as a limitation for peach cultivation in the northern basin of the Mediterranean. This is consistent with our results indicating that chilling requirement was systematically satisfied for the reference period 1996-2015 in the entire French territory between mid-December and mid-February, with the milder regions satisfying it later. On the other hand, frost damage and insufficient heat accumulation are known to be primary limitations to the cultivation in Europe of peaches and other *Prunus* cultivars (e.g. apricot and cherry, see Julian et al., 2007; Reig et al., 2013; Chmielewski et al., 2017; Vitasse and Rebetez, 2018). For peaches, this was reflected in our study for those areas outside the current French peach production regions, such as the north-western areas of Occitanie and the alpine part of Auvergne Rhône-Alpes. Note that even if an area is suitable for the cultivation of very few cultivars, generally such an area cannot be considered cost-effective for peach production because the product would be available only for a limited time and late in the season (Layne & Bassi, 2008). Our model identifies the locations of Occitanie, Auvergne-Rhône-Alpes (ARA) and Provence-Alpes-Côte d’Azur (PACA) as the most suitable area for peach cultivation in France. This outcome of our model is consistent with the fact that they currently provide more than 90% of the France’s total peach production (Talpin, 1954; Ministère de l’Agriculture et de l’Alimentation, 2019).

### 4.2 Peach thermal niche in the future

Frost damage and heat accumulation constraints are predicted to wane in northern regions of continental France, namely in Pays de la Loire, Centre-Val de Loire and Bourgogne-France-Comté by the end of the 21^st^ century, a result that is in line with projections of the northward expansion of other cultivars due to global warming (Chmielewski & Rötzer, 2001). If warmer winter temperatures determining earlier blooming are not followed by warmer spring temperatures at blooming time, an higher risk of spring frost can occur (Liu et al., 2018; Vitasse & Rebetez, 2018). On the other hand, such risk can be neglected when also spring temperatures increase below the frosting threshold (Eccel et al., 2009; Chmielewski et al., 2017). Our results suggest that, due to the specific interplay between peach phenology and French predicted climate change the second situation will likely occur in France.

According to our findings, the decline in winter chill will become the major limitation for peach cultivation. Blooming failure due to mild winters was first observed in northern Africa (Ghrab et al., 2014) and it is likely to become the norm in southern France by the end of the century. The limitation of the production area for peach appears to be even more severe using our model under the RCP 8.5 scenario (Figure S6), with a further shift of the suitable area towards the alpine and northern continental regions of the country. Our results are consistent with the growing concern regarding the impact of global warming on the endodormancy phenological phase. Several authors already provided evidence of abnormal patterns of bud break and fruit development in Europe (Legave et al., 1983; Erez & Couvillon, 1987; Viti et al., 2010) and relevant economic issues (Jackson & Hamer, 1980; Baldocchi & Wong, 2008; Luedeling et al., 2009; Ghrab et al., 2014).

Beyond shifting the geographies of the thermal niche of peaches, climate change is expected to alter peach phenology and physiology. We predict that blooming will occur between 5 and 20 days earlier than usual by the end of the century, depending on the area and the emission scenario. Earlier blooming has also been predicted by other authors for other species (Chmielewski et al., 2004; Dose & Menzel, 2004; Menzel et al., 2006; Estrella et al., 2007; Primack et al., 2009; Fujisawa & Kobayashi, 2010; Jochner et al., 2016; Parker & Abatzoglou, 2017). When dealing with commercial species, it is worth noting that faster fruit ripening due to increased temperatures affects fruit growth physiology (Lescourret & Genard, 2005). This might impair the quality, in terms of organoleptic properties, of the final product (Peiris et al., 1996).

Needless to say, all predicted changes in peach phenology and thermal niche depend on methodological choices regarding both the climate change scenarios and the phenological models. Reliable models to estimate blooming dates are still lacking due to the complex interplay of the processes governing endo- and ecodormancy breaks (Bartolini et al., 2018) and discrepancies between different models are unfortunately still the norm rather than the exception (Chmielewski et al., 2012; Andreini et al., 2014). The main challenge is to achieve a better understanding of the chill accumulation and endodormancy break processes (Luedeling, 2012). New empirical data that provide measures of endodormancy break dates, such as those studied by Chuine et al. (2016), are to our view urgently needed so as to conceive, calibrate and validate models aimed at testing further hypotheses. In our model we assumed that plant water needs would be met through irrigation. However, climate changes are expected to increase the frequency of both drought and heavy precipitation events (IPCC, 2013), factors that surely will impact water availability for agriculture (Elliott et al., 2014). Moreover, shifts in the plant phenological dates and geographical range may result in another possible mismatch with pollinator phenology (Scaven & Rafferty, 2013) and photoperiod (Hänninen & Tanino, 2011; Way & Montgomery, 2015). This latter will not change as the climate warms, yet the potential break of the synchrony in the plant-climate integrated system remains highly probable. Likewise, plants may be exposed to new parasites and consequent diseases (Chakraborty & Newton, 2011). Only when a transdisciplinary and broader approach will provide further evidence, the inclusion of these processes, and possibly others, in our framework will be possible.

### 4.3 Adapting to climate change

Despite the uncertainty which is inherent in any projection, there is no doubt that climate changes will affect peach phenology and the geographical range of the thermal niche. Our study corroborate the intuition that current suitable areas can turn into unsuitable and viceversa. This will occur to the extent that adaptation strategies will be required. As insufficient winter chill will be the major issue, some research breeding studies have been started to search for cultivars with low or no chilling requirements. Such cultivars are already being utilized in tropical and subtropical climates (Layne & Bassi, 2008). However, chill requirements evolved to protect plants from frost damages and “artificially” lowering them might increase the impacts of frost damages. Some chemicals have been shown to be effective in forcing flowering and bud burst in dormant plants (Erez et al., 2008; Ashebir et al., 2010). However, they are effective only under given environmental conditions (Campoy et al., 2011) and they make use of phytotoxic compounds (Luedeling, 2012). It has also been shown that winter agronomic practices (like sprinkling water over shoots) can reduce the buds temperature by evaporative cooling (Erez, 1995).

An alternative adaptation strategy could consist in moving the peach production into new suitable areas. This strategy has three obvious drawbacks: *i*) the costs of crop translocation is substantial, *ii*) additional costs can be incurred in transporting fruit to centralized processing and distribution facilities (Parker & Abatzoglou, 2017), and *iii*) the sociological implications of the growers’ behavior should be considered as they will ultimately govern the reality of peach cultivation in new regions.

## 5 Conclusions

Our study provides a model to produce data-informed maps that quantitatively allow the analysis of the geographical limitations and challenges that the peach industry may face under climate change in France. Here, we used the best currently available information about phenology of fruit trees. We developed a novel phenological model that is not site specific, thus it can be applied to other systems where climate change scenarios are available. It is clear that global warming will impact the economic viability of peach production and models can support actions to be taken now to adapt to change.

## Acknowledgments

This work has been partly carried out as part of the INRA Metaprogramme ACCAF (project 429 CLIF). We thank Agroclim unit (INRAE) for providing data from the different weather stations via their web service Climatik, and the TEMPO French Network (https://tempo.pheno.fr/) for providing the access to different phenological data resources in their portal. This work has been carried out as part of the ECOVERGER project. This action is led by the Ministry for Agriculture and Food and the Ministry for an Ecological and Solidary Transition, with the financial support of the French Biodiversity Agency on “Resistance and Pesticides” research call, with the fees for diffuse pollution coming from the Ecophyto plan.

## Supporting Information

**Figure S1:**
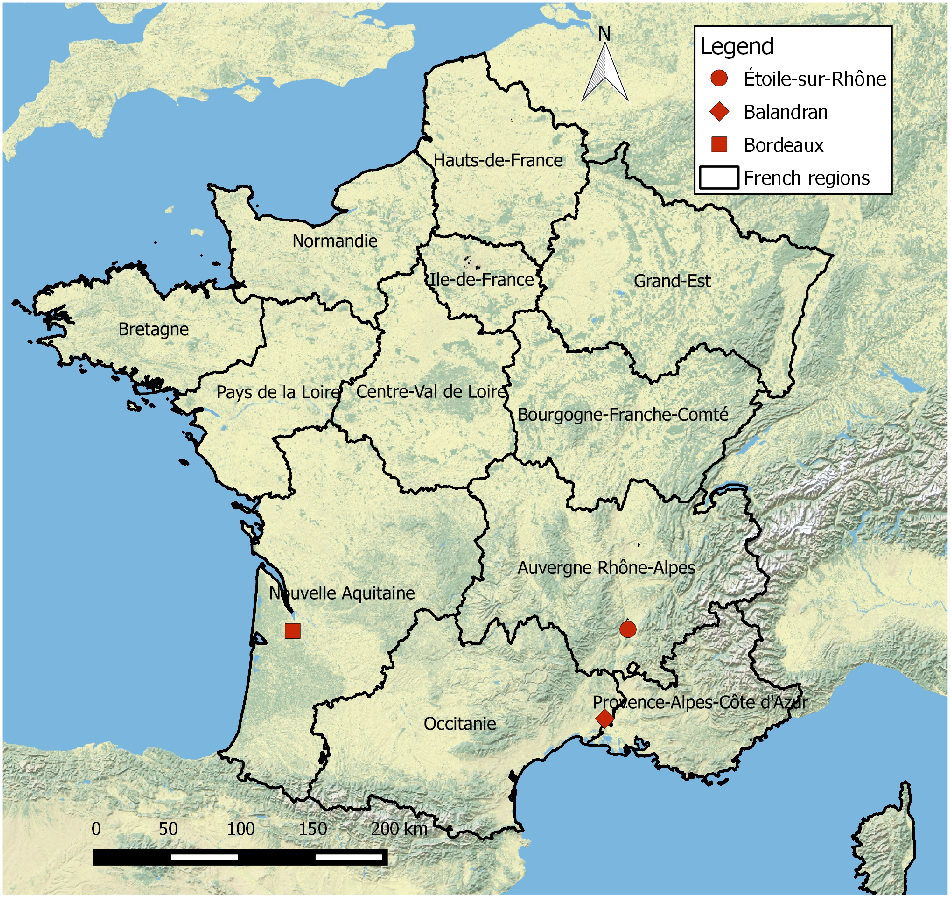
French continental regions and sampling sites of blooming and ripening dates.

**Table S1:** Meteorological stations. Meteorological station coordinates used for the period of interest in Balandran, Bordeaux and Étoile-sur-Rhone sites

| Phenological data site | Years of interest | Longitude | Latitude |
| --- | --- | --- | --- |
| Balandran | 1986-2004 | 4° 26' E | 43° 47' N |
|  | 2005-2008 | 4° 28' E | 43° 45' N |
| Bordeaux | 1986-1999 | 0° 35' W | 44° 47' N |
| Étoile-sur-Rhône | 1996-2005 | 4° 44' E | 44° 35' N |

**Figure S2:**
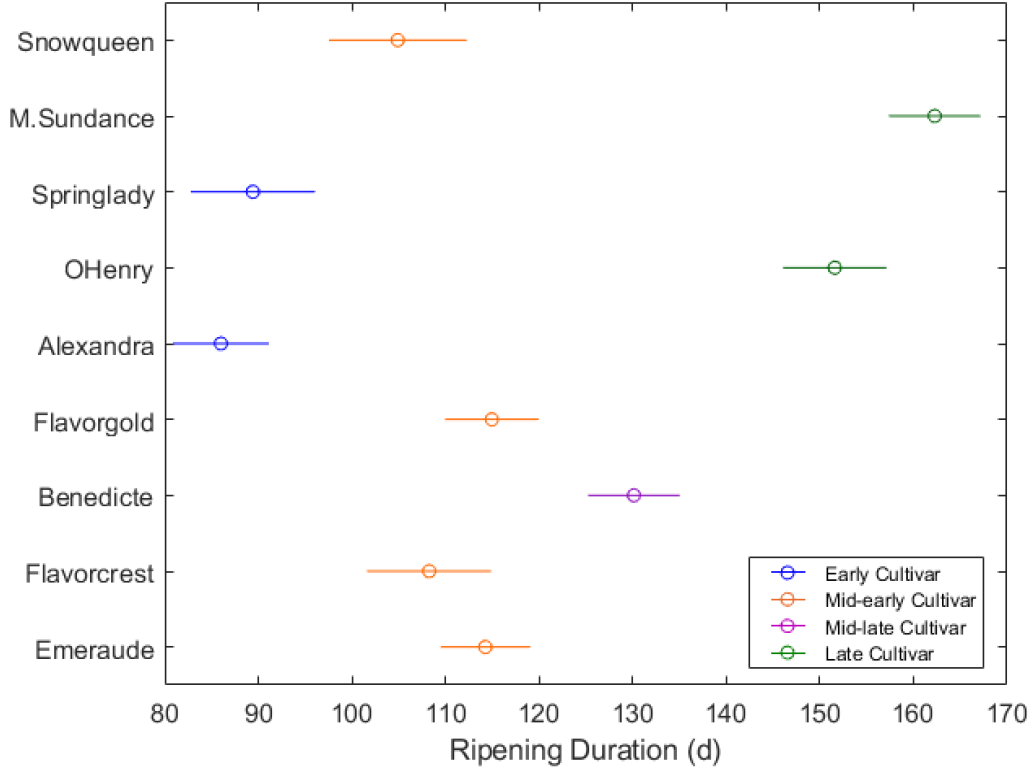
Peach cultivar and ripening duration. Cultivar clustering of the ripening duration period (multiple t-test, Scheffé’s procedure). In blue, orange, purple and green are represented early, mid-early, mid-late and late cultivars, respectively.

**Figure S3:**
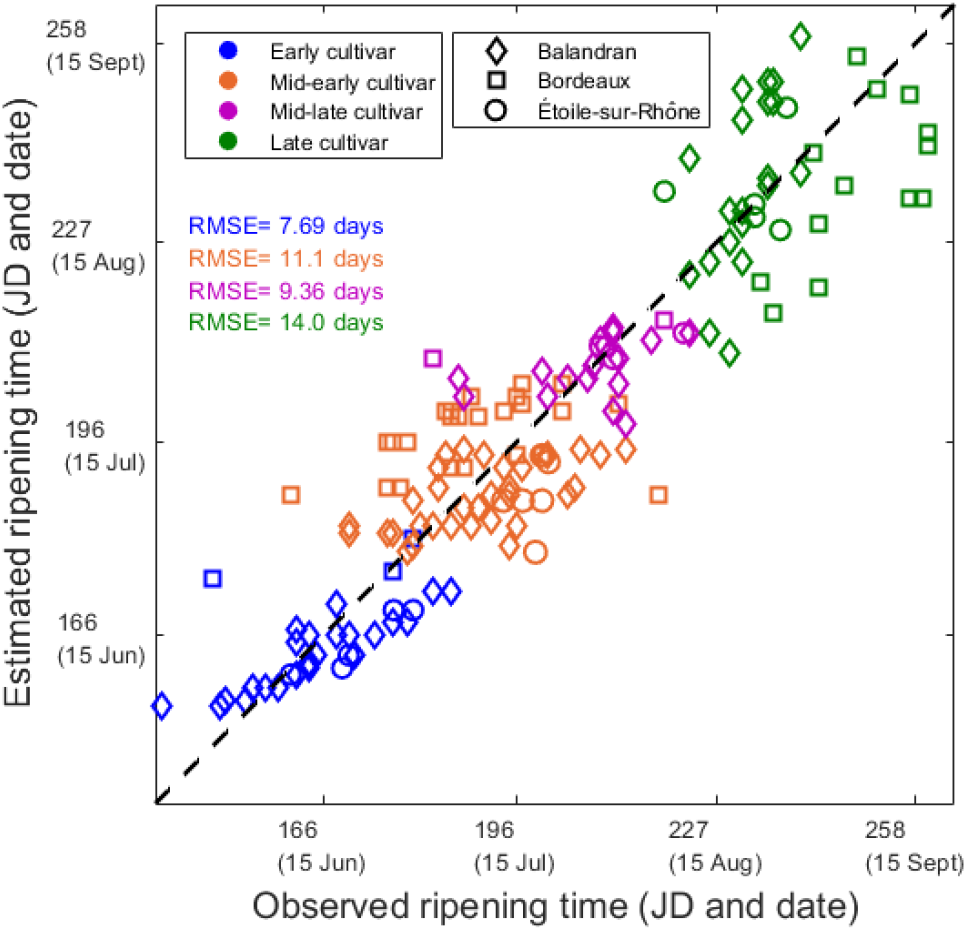
Observed and predicted dates of ripening. The ripening date has been predicted by summing the predicted ripening duration to the predicted (not observed as in fig. 1) date of blooming. Diamonds, squares and circles refer to the sites of Balandran, Bordeaux and Étoile-sur-Rhône, respectively. Ripening date is represented in blue for early, in orange for mid-early, in purple for mid-late and in green for late cultivars.

**Figure S4:**
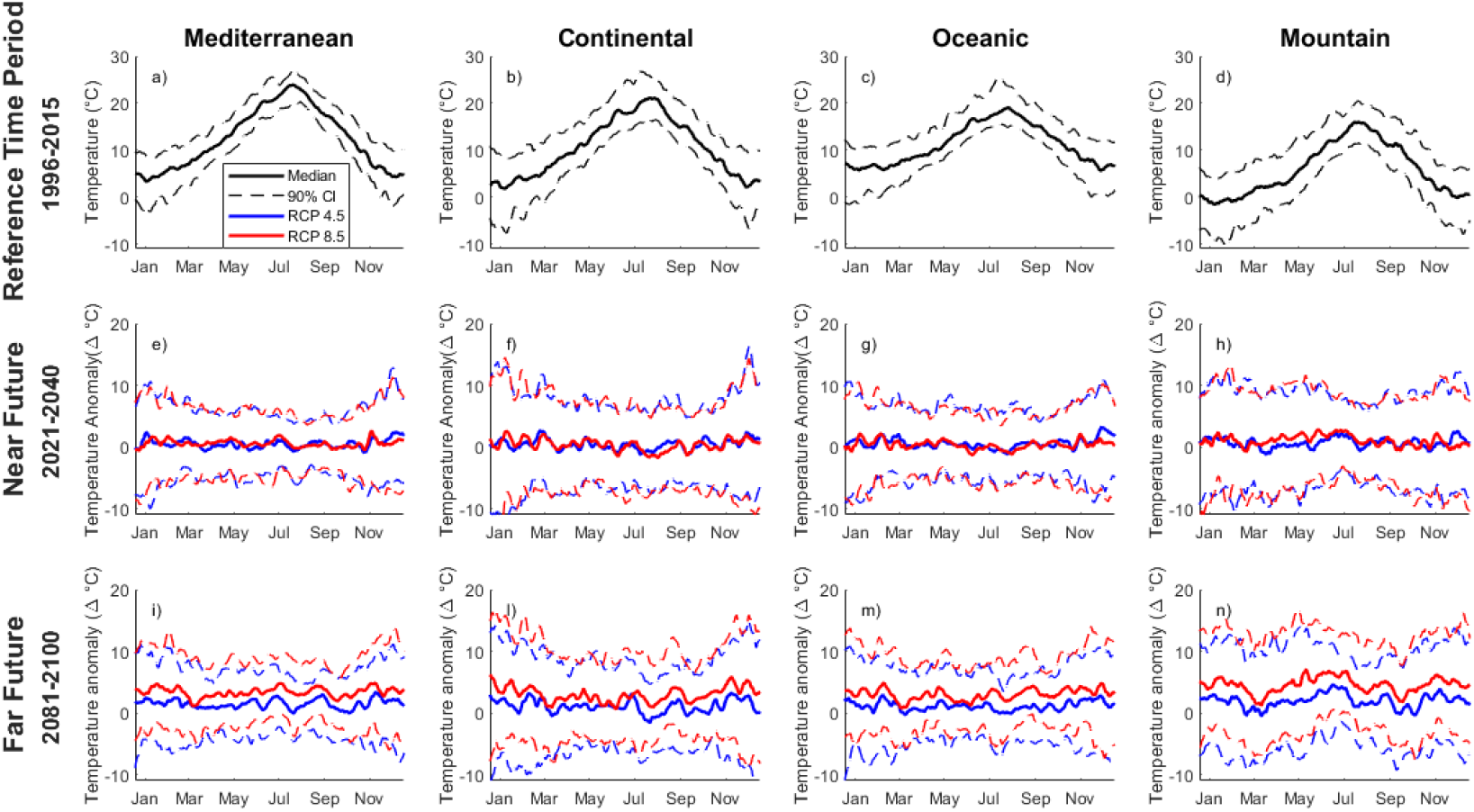
Temperatures in Mediterrean, continental, oceanic and mountain areas of the French territory: hindcast and forecast. Mean daily temperature, averaged with a moving window of 10 days, in the reference period (1996-2015) in black (a, b, c, d) and relevant anomalies in the near (2021-2040) and far (2081-2100) future in four areas of 24 × 24 km^2^ representative of different climatic zones: Mediterrean (43° 53′ N 5° 4′ E), continental (48° 48′ N 4° 14′ E), oceanic (48° 11′ N 2° 59′ W) and mountain (42° 53′ N 0° 2 E). In black the temperatures in the reference period, in red and blue the temperature anomalies, compared to the reference period, in RCP 4.5 and in RCP 8.5, respectively. Solid lines represent the median daily temperatures, while dashed ones the 5% and 95% percentile.

**Table S2:** Seasonal temperature anomaly in Mediterrean, continental, oceanic and mountain areas of the French territory: forecast. Average seasonal temperature (°C) in the reference period (1996-2015) and temperature anomaly (Δ° C) in the near (2021-2040) and far (2081-2100) future, in four different French climatic zones (Mediterranean, continental, oceanic and mountain). Relative 90% confidence interval are reported in square brackets. See Figure S4 for details about the climatic areas.

| Average Temperature ($^{\circ}\text{C}$) | | | | | |
| --- | --- | --- | --- | --- | --- |
| Time period | Season | Mediterranean | Continental | Oceanic | Mountain |
| Reference | Spring | 11.2 [12.3;14.7] | 11.6 [10.2;12.3] | 11.3 [9.9;12.0] | 5.4 [3.5;5.1] |
|  | Summer | 21.1 [19.6;22.2] | 18.6 [17.1;21.0] | 17.4 [16.0;19.2] | 13.5 [11.7;15.8] |
|  | Autumn | 10.0 [8.7;11.3] | 8.1 [6.3;9.7] | 9.8 [8.3;11.5] | 4.8 [2.7;6.2] |
|  | Winter | 5.2 [3.3;6.8] | 3.8 [0.83;5.35] | 6.5 [4.3;7.3] | -0.24 [-2.4;1.2] |
| RCP 4.5-Temperature Anomaly ($\Delta^{\circ}\text{C}$) | | | | | |
| Time period | Season | Mediterranean | Continental | Oceanic | Mountain |
| Near future | Spring | -0.071 [-2.3;2.7] | 0.22 [-2.4;2.2] | 0.23 [-2.0;2.2] | 0.044 [-3.0;5.2] |
|  | Summer | 0.44 [-1.5;2.2] | 0.39 [-2.3;2.2] | 0.052 [-1.4;2.4] | 1.2 [-1.6;3.5] |
|  | Autumn | 0.39 [-2.0;3.2] | -0.045 [-1.2;3.8] | 0.52 [-0.93;3.5] | 0.015 [-2.4;5.0] |
|  | Winter | 1.3 [-2.2;3.1] | 1.2 [-3.0;4.3] | 0.93 [-1.3;3.2] | 1.5 [-1.6;3.4] |
| Far future | Spring | 1.2 [-0.78;0.87] | 0.70 [-1.6;3.7] | 0.92 [-1.3;3.6] | 2.0 [-1.5;6.5] |
|  | Summer | 1.4 [0.36;3.1] | 0.51 [-2.24;3.1] | 0.75 [-0.70;2.4] | 2.5 [0.47;4.3] |
|  | Autumn | 1.4 [-1.1;3.2] | 1.6 [-1.3;4.0] | 1.6 [0.94;4.0] | 1.7 [-1.3;5.1] |
|  | Winter | 2.1 [-1.0;4.6] | 1.3 [-2.6;5.1] | 1.1 [-2.0;4.2] | 2.2 [-1.0;5.1] |
| RCP 8.5-Temperature Anomaly ($\Delta^{\circ}\text{C}$) | | | | | |
| Time period | Season | Mediterranean | Continental | Oceanic | Mountain |
| Near future | Spring | 0.39 [-2.3;2.2] | 0.35 [-1.8;3.1] | 0.18 [-1.5;2.2] | 1.4 [-1.9;4.9] |
|  | Summer | 0.21 [-1.9;2.5] | -0.28 [-4.4;3.0] | -0.51 [-2.6;2.2] | 1.4 [-1.2;3.6] |
|  | Autumn | 0.22 [-0.99;2.3] | 0.42 [-4.4;2.6] | 0.33 [-1.6;2.7] | 0.30 [-2.4;3.8] |
|  | Winter | 1.0 [-2.1;3.1] | 0.71 [-2.9;4.2] | 0.78 [-2.0;2.4] | 0.83 [-1.6;2.9] |
| Far future | Spring | 2.63 [1.2;5.6] | 1.7 [0.41;5.3] | 1.8 [0.94;4.7] | 4.6 [0.84;9.1] |
|  | Summer | 3.6 [2.0;5.9] | 2.9 [-0.41;5.3] | 3.0 [0.47;5.6] | 5.1 [3.1;7.5] |
|  | Autumn | 3.3 [1.4;5.4] | 3.50 [0.66;6.0] | 3.4 [1.2;5.9] | 4.2 [1.7;6.6] |
|  | Winter | 3.2 [1.2;6.1] | 3.04 [1.3;8.3] | 2.8 [2.0;7.1] | 4.1 [1.6;7.6] |

**Figure S5:**
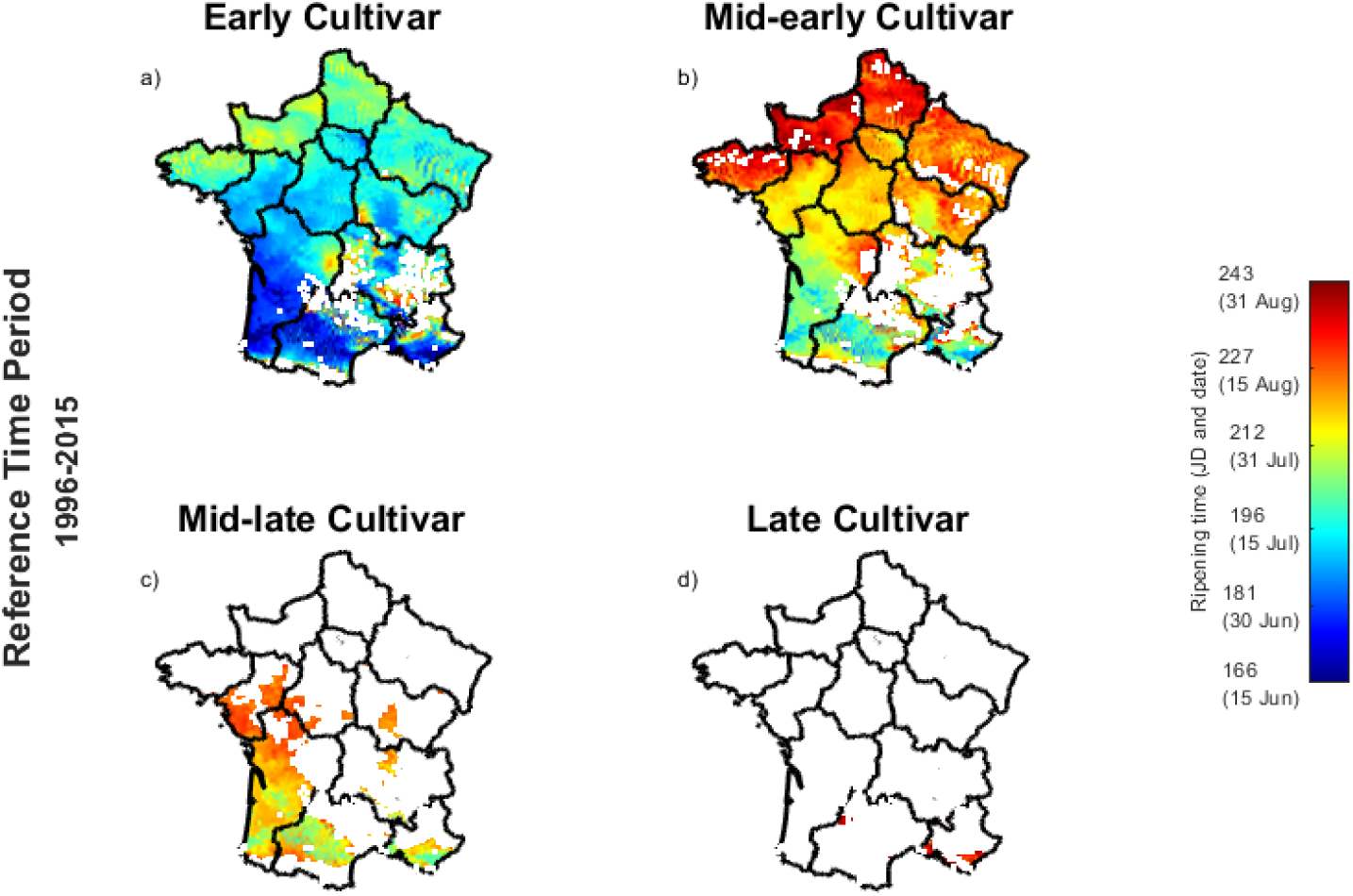
Peach cultivar ripening time in the French continental regions in the reference period (1996-2015). Predicted ripening date, in Julian Days (JD) and calendar dates (within brackets), in the reference time period (1996-2015) for (a) early, (b) mid-early, (c) mid-late and (d) late cultivars. White cells represent those areas that are not suitable for the cultivar production. For each cell (8× 8 km^2^), the average value over the considered 20 years is reported.

**Figure S6:**
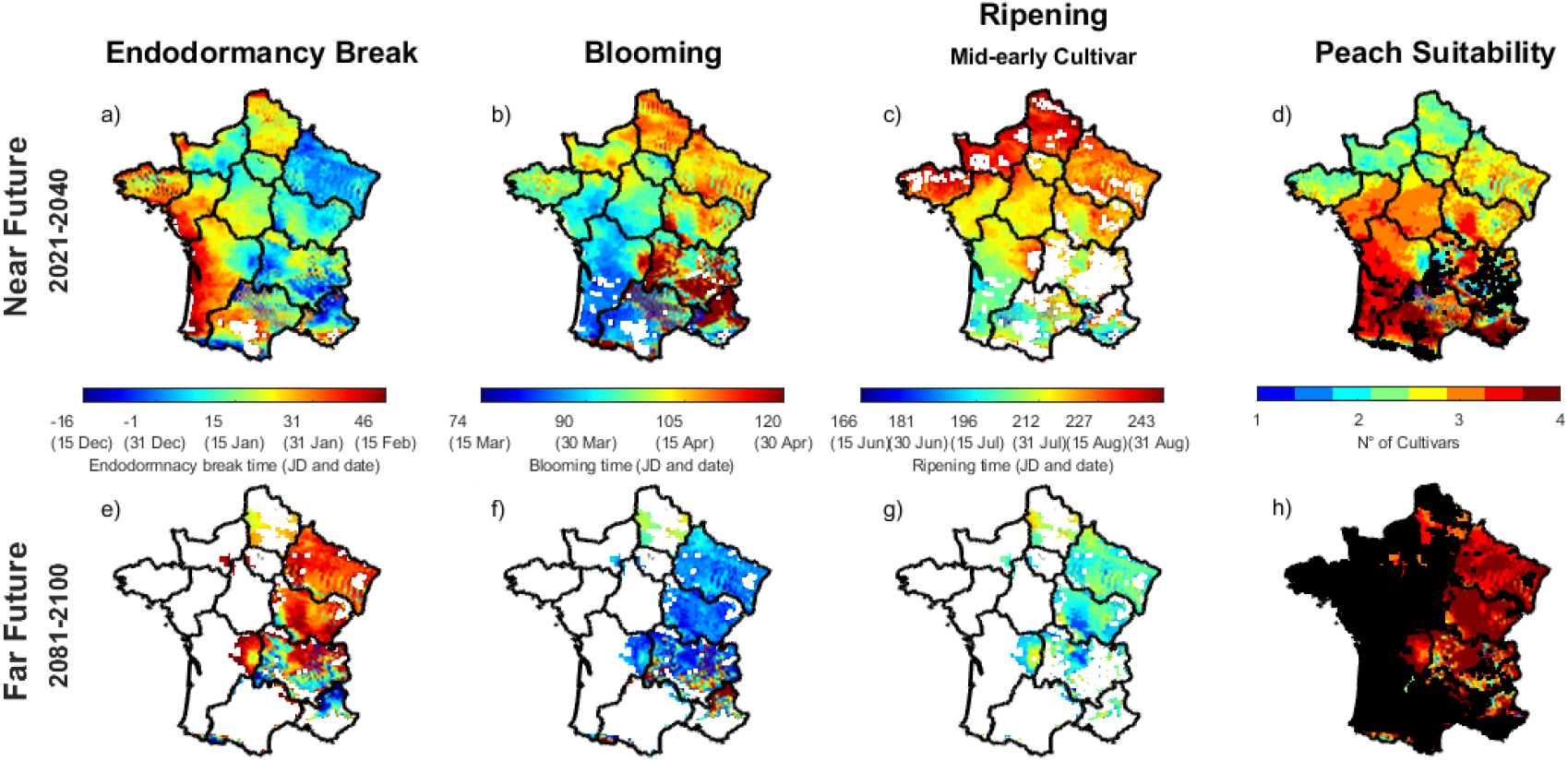
Peach phenological times and thermal niche for cultivation: scenarios forcing our model with RCP 8.5 in the French continental regions for the near (2021-2040, upper panels) and far (2081-2100, lower panels) future. (a,e) Endodormancy break, (b,f) blooming, (c,g) ripening of the mid-early cultivars dates, in Julian Days (JD) and calendar dates (within brackets), and (d,h) peach suitability. For each map cell (8×8 km^2^), the average value over the considered 20 years is reported. White map cells (a-c and e-g) represent those areas where the phenological event did not occur for at least two years. Likewise, black cells (d and h) represent those areas where no peach cultivar could be cultivated for at least two years (out of 20).

